# Feedback control of the heat shock response by spatiotemporal regulation of Hsp70

**DOI:** 10.1101/2024.01.09.574867

**Authors:** Rania Garde, Annisa Dea, Madeline F. Herwig, David Pincus

## Abstract

Cells maintain homeostasis via dynamic regulation of stress response pathways. Stress pathways transiently induce response regulons via negative feedback loops, but the extent to which individual genes provide feedback has not been comprehensively measured for any pathway. Here, we disrupted induction of each gene in the *Saccharomyces cerevisiae* heat shock response (HSR) and quantified cell growth and HSR dynamics following heat shock. The screen revealed a core feedback loop governing expression of the chaperone Hsp70 reinforced by an auxiliary feedback loop controlling Hsp70 subcellular localization. Mathematical modeling and live imaging demonstrated that multiple HSR targets converge to promote Hsp70 nuclear localization via its release from cytosolic condensates. Following ethanol stress, a distinct set of factors similarly converged on Hsp70, suggesting that nonredundant subsets of the HSR regulon confer feedback under different conditions. Flexible spatiotemporal feedback loops may broadly organize stress response regulons and expand their adaptive capacity.

## INTRODUCTION

Stress response pathways enable cells to adapt to environmental changes and survive. Stress responses deactivate processes that are no longer adaptive and induce new gene expression programs for survival and growth in the new conditions. However, to avoid overshooting adaptation, the stress response must also efficiently turn off. It is unclear how response dynamics are tuned by the induction of individual downstream target genes. How and to what extent do stress responses integrate feedback from their suite of effectors?

The heat shock response (HSR) is an ancient, conserved, and prototypical stress response pathway under the control of the master regulator Hsf1 in eukaryotes (*1, 2*). When an environmental stressor or internal dysfunction causes an excess of newly synthesized/unfolded, misfolded or mis-targeted proteins to accumulate in the cytosol or nucleus, Hsf1 transcriptionally induces molecular chaperones and other factors involved in protein folding, disaggregation, and degradation (*3, 4*). Once protein homeostasis (proteostasis) is restored, molecular chaperones become available to bind and deactivate Hsf1. Previously, the heat shock-induced, Hsf1-dependent transcriptome was characterized in *S. cerevisiae* using genomic and chemical genetic methods, revealing a compact set of 42 target genes that both show increased Hsf1 occupancy in their enhancer region and are dependent on Hsf1 for their transcription during heat shock (*5*). Here we ask how the transcriptional induction of these 42 individual genes – collectively referred to as the HSR regulon – determine the dynamics of Hsf1 activity.

In addition to addressing basic questions of adaptive regulation of stress response pathways, understanding how cells dynamically control Hsf1 activity is relevant to human health. Indeed, Hsf1 mis-regulation in both directions – either too much or too little activity – is associated with disease. In aggressive cancers, Hsf1 is often hyperactivated or overexpressed due to Hsf1 locus amplification (*6*). This increased Hsf1 activity not only induces Hsp90 and other chaperones to support folding of oncoproteins but also drives a cancer-specific gene expression program that supports malignancy both in the tumor cells and the supporting microenvironment (*7–11*). On the other hand, in neurodegenerative disorders, aggregates of proteins remain unresolved and are thought to sequester chaperones, hampering general cellular processes and triggering further protein aggregation (*12, 13*). As such, loss of Hsf1 has been implicated in Huntington’s disease and increasing Hsf1 activity has been proposed as a therapeutic avenue for neurodegenerative disease with broad potential (*14–16*). Thus, resolving how Hsf1 activity is tuned in healthy cells may inform these disease mechanisms.

Mechanistically, Hsf1 is regulated by an Hsp70-based negative feedback loop (*17, 18*). The chaperone Hsp70 directly binds and represses Hsf1 in the nucleoplasm under non-stress conditions (*19, 20*). Upon heat shock, Hsp70 dissociates from Hsf1 and is targeted to protein condensates in the cytoplasm and nucleolar periphery via its cofactor, the J-domain protein Sis1 (*21, 22*). This leaves Hsf1 free to itself form active transcriptional condensates and induce its target genes, including multiple Hsp70 paralogs (*23, 24*). Induction of Hsp70 is required for Hsf1 deactivation, and within fifteen minutes of heat shock, Hsf1 is rebound by Hsp70 and Hsf1 transcriptional activity is repressed again (*17, 20*). Thus, Hsf1 is dynamically tuned via its direct interactions with Hsp70. While these precedents were established in yeast, the same mechanisms have been shown to be largely conserved in mammalian cells (*25–27*).

Immediately upon heat shock, prior to or coincident with induction of the HSR regulon, the translation initiation machinery and mRNAs form reversible condensates known as stress granules (*28–31*). Stress granules are regulated by chaperones including Hsp70, J-domain proteins, Hsp104, small heat shock proteins, and Hsp90—all of which are Hsf1 targets (*32–34*). In addition to protein-RNA condensates, protein-only condensates and secretory vesicles also recruit these same chaperones during heat shock (*35–40*). Therefore, though Hsp70-based negative regulation is the only direct Hsf1 regulation known, other targets may influence Hsf1 directly or indirectly by regulating localization of the chaperone machinery to the various stress-induced condensates sub-localized to regions of the cytosol or nucleus.

Here, we collected >10^7^ single cell fluorescence measurements and >10^4^ growth measurements to comprehensively dissect feedback regulation in the HSR. First, we characterized the transcriptional dynamics of each HSR target gene during heat stress in *S. cerevisiae*. Next, we disrupted the transcriptional induction of each target gene by deleting the 9-25 bp Hsf1 binding region in the upstream regulatory region via CRISPR/Cas9-mediated genome editing and observed how impaired induction of each target influences global output of the HSR and growth at elevated temperature. Additionally, we repeated the regulon-wide screen in response to ethanol rather than heat shock. Finally, using mathematical modeling and live cell imaging, we demonstrate that that the feedback architecture of the HSR is remarkably simple: a core feedback loop controlling the expression of Hsp70 is reinforced by an auxiliary feedback loop – comprised of condition-specific subsets of the HSR regulon – controlling the interaction of Hsp70 with cytosolic condensates and thereby regulating Hsp70 nuclear localization. Such a flexible feedback hierarchy that converges to control both expression and subcellular localization of key effectors may broadly characterize stress response regulons.

## RESULTS

### Regulon-wide measurement of HSR gene expression dynamics

Previously, nascent transcript sequencing coupled to Hsf1 depletion and genome-wide Hsf1 ChIP-seq revealed that 42 target genes are directly bound by Hsf1 and dependent on Hsf1 for their transcription upon heat shock (Figure 1A) (*5*). To establish the transcriptional dynamics of the HSR regulon at single cell resolution, we measured the expression of each Hsf1 target gene in 10^4^ single cells at 10 time points over a four-hour heat shock time course. To this end, we generated a library in which we tagged each Hsf1 target gene in the genome at the 3′ end with a P2A ribosome skip sequence followed by mScarlet (Figure 1B). The polycistronic mRNAs expressed in the 42 reporter strains all have the same 3′ untranslated regions, so differences in fluorescence signal should reflect differences in transcription more than mRNA stability.

**Figure 1.**
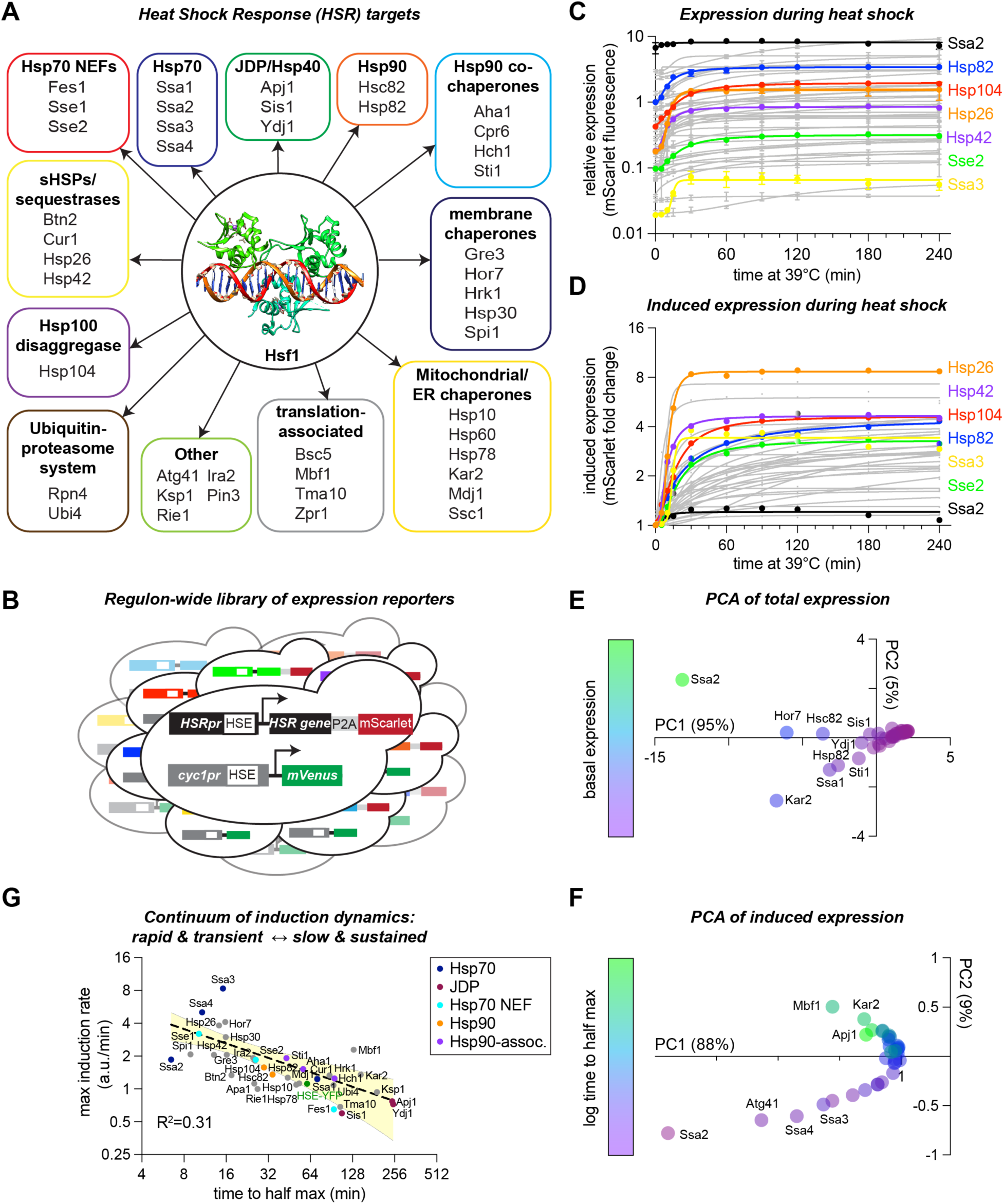
Regulon-wide measurement of HSR gene expression dynamics. **A)** 42 genes are directly bound by Hsf1 and dependent on Hsf1 for their transcription. **B)** Schematic of gene expression reporter strategy. P2A-mscarlet is fused to each gene in the HSR regulon for measurement in single cells. **C)** mScarlet levels over a heat shock time course for all Hsf1 target genes, measured by flow cytometry. Each data point represents the mean of three biological replicates, connecting lines represent the non-linear curve fit for each time trace. **D)** Data from (C) replotted to highlight fold change mScarlet lduring heat shock. **E)** Principal component analysis (PCA) of the time course data in (C) showing coordinates of each gene in the first two PCs and color coded by expression level. **F)** PCA of the normalized data in (D) showing coordinates of each gene in the first two PCs and color coded by the log of the time to half max. **G)** Maximum induction rate plotted verses time to half maximum expression calculated with parameter estimates from the non linear curve fits of induced expression. Yellow shaded area represents 95% CI on the bounds of the fit.

Consistent with previous RNA-seq experiments (*4, 5, 21*), the expression level of Hsf1 target genes varies by three orders of magnitude across the regulon (Figure 1C). Regulon-wide basal expression level measured by mScarlet correlates well with transcript measured by sequencing (r = 0.82, Figure S1A). Upon heat shock, all target genes were induced over the time course with magnitudes ranging from <10% to >8-fold (Figure 1D, Figure S1B).

We applied principal component analysis (PCA) to identify modes of variation across the dataset. Remarkably, 95% of the total variance was explained by the first principal component. Given the three orders of magnitude range expression, we found PC1 to be associated with basal mScarlet expression levels (Figure 1E). Previously, we showed that basal expression across the regulon is determined by a combination of the biochemical affinity of Hsf1 for each binding site and the accessibility of the binding site in the chromatin landscape (*5*), so we already have insight into the molecular basis for this mode of variation. To focus on the variation in the dynamics, we performed PCA on the induced expression dataset (basal level-normalized). Plotting the resulting PC1 against PC2 reveals a semi-continuous manifold that corresponds to the time it takes for each gene to reach its half-maximum induction (Figure 1F). Upon further analysis, we observed an apparent tradeoff between the time to half-max induction and the maximum rate of induction (Figure 1G). These data imply that there is a continuum of gene expression profiles driven by Hsf1, ranging from genes that turn on rapidly, robustly, and transiently to those that turn on slowly, weakly, but in a sustained fashion.

### Regulon-wide screen for HSR feedback regulators during heat shock

To determine which Hsf1 target genes are feedback regulators of the HSR, we created two additional regulon-wide libraries by deleting the empirically determined Hsf1 binding sites – known as heat shock elements (HSEs) – in the endogenous regulatory region of each target gene with scarless CRISPR/Cas9-mediated genome editing (see methods) (Figure 2A) (*5*). First, we generated HSE deletion (ΔHSE) strains in the mScarlet reporter library used above and quantified the extent to which each ΔHSE mutation alters expression of the linked gene throughout the heat shock time course. We successfully generated ΔHSE strains for 39/42 genes in the HSR regulon. We were unable to obtain HSE deletions in the regulator regions of *AHA1*, *STI1*, or *KSP1*, likely for technical rather than biological reasons. Among the set of Hsf1 target genes, *HSP26* is unique in having two distinct clusters of HSEs rather than a single Hsf1 binding peak. Simultaneous disruption of both sites nearly abrogated mScarlet induction, while individual disruption of each of the sites resulted in differential induction dynamics, separately impairing rapid and sustained induction (Figure 2B). Across the HSR regulon, HSE deletion reduced induction during heat shock for all genes except *HSP30*, *HOR7*, and *YDJ1* (Figure 2C). Residual heat shock-induced expression of these genes may be due to undefined *cis*-elements and/or transcription factors. Basal expression was also reduced for many genes, consistent with prior reports that Hsf1 drives constitutive expression of a subset of its target genes (*4, 41*).

**Figure 2.**
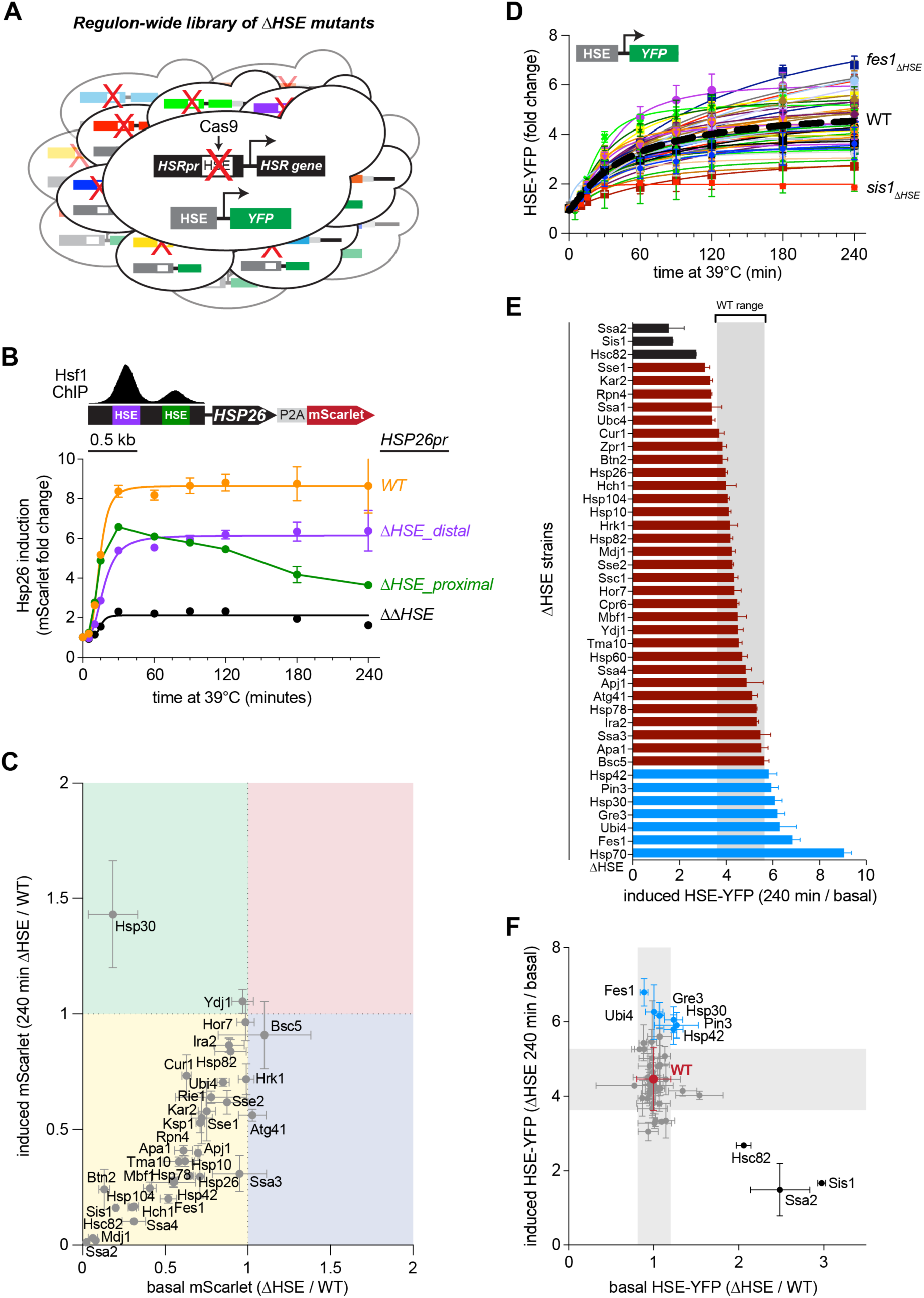
Regulon-wide screen for HSR feedback regulators during heat shock. **A)** Schematic of the library of ΔHSE mutants expressing the synthetic reporter of the HSR (HSE-YFP). **B)** Effect of single and double disruption of the HSEs in the *HSP26* promoter on the expression of P2A-mScarlet reporter over a heat shock time course. **C)** mScarlet levels in each ΔHSE mutant relative to its wild type counterpart under plotted in basal conditions versus four hours of heat shock. Each data point represents the mean and standard deviation of three biological replicates for both ΔHSE strain and wild type counterpart. **D)** HSE-YFP fold change over a heat shock time course for all ΔHSE mutants, measured by flow cytometry. Dashed black line represents WT average. Each data point represents the mean and standard deviation of three biological replicates. **E)** HSE-YFP fold change after four hours of heat shock for each ΔHSE mutant and the Hsp70ΔFBL mutant. WT range in gray represents the range of HSE-YFP levels across 20 biological replicates of WT. Bars show the mean of 3 biological replicates, error bars the standard deviation. **F)** HSE-YFP level after four hours of heat shock versus levels basal conditions for each ΔHSE mutant. Each data point represents the mean and standard deviation of three biological replicates. WT range in gray represents the mean and standard deviation of 20 biological replicates.

Next, we constructed a second ΔHSE strain library in which each gene retains its endogenous 3′ UTR to test the consequences of disrupting transcriptional induction without perturbing mRNA stability. The library also harbors a synthetic reporter of Hsf1 activity (HSE-YFP) integrated into the genome to serve as a standardized measurement of HSR activity in single cells (*19, 42*). As with the mScarlet reporters, we measured HSE-YFP levels in the ΔHSE library over four hours of heat shock in individual cells by flow cytometry (Figure 2D). HSE-YFP levels varied four-fold across the mutants, but most mutants had HSE-YFP values after four-hour heat shock that fall within the wild type reproducibility range, suggesting – with the caveat that many ΔHSE mutants show incomplete loss of expression – that the induction of most Hsf1 targets neither directly regulates nor indirectly affects Hsf1 activity during heat shock (Figure 2E, Figure S2A).

We previously engineered a strain, Hsp70ΔFBL, lacking Hsf1-dependent induction of the four genes encoding cytosolic Hsp70 in yeast (*ssa1/2/3/4*) that showed a pronounced defect in deactivating the HSE-YFP reporter following sustained heat shock (*17*). None of the ΔHSE mutants approached the increased HSE-YFP levels we observed in Hsp70ΔFBL (Figure 2E). However, six ΔHSE mutants showed significantly increased, and three showed significantly reduced, HSE-YFP levels compared to wild type after four hours of heat shock (p < 0.05, two-tailed t-test). The three mutants with reduced induction, *ssa2ΔHSE, sis1ΔHSE,* and *hsc82ΔHSE*, all had increased basal HSE-YFP levels (Figure 2F, Figure S2B). The increased basal levels account for their reduced induction, consistent with previous reports that these factors function as basal repressors (*43, 44*), indicating that these are not positive feedback regulators. By contrast, except for *hsp30ΔHSE* in which the mScarlet reporter is still induced during heat shock (Figure 2C), the ΔHSE mutants with elevated HSE-YFP levels after four hours of heat shock – *fes1ΔHSE*, *ubi4ΔHSE*, *gre3ΔHSE*, *pin3ΔHSE*, and *hsp42ΔHSE* – are candidate negative feedback mutants.

### Assessment of functional redundancy among key chaperone families

The three mutants with elevated HSE-YFP levels under basal conditions, *ssa2ΔHSE, hsc82ΔHSE*, and *sis1ΔHSE,* encode members of the cytosolic Hsp70, Hsp90 and JDP chaperone families, respectively. In the HSR regulon, three additional genes encode cytosolic Hsp70 (*SSA1*, *SSA3*, and *SSA4*), one additional gene encodes Hsp90 (*HSP82*), and two additional genes encode JDPs (*APJ1* and *YDJ1*). Chaperones of each family may have redundant functions that can be revealed with multiple mutations; Hsp70 provides a demonstrative case. The relative expression level and induction dynamics of each Hsp70 paralog as measured by mScarlet levels span the full range of the library with the levels of Ssa3 < Ssa4 < Ssa1 < Ssa2 (Figure 3A, top). While none of the single ΔHSE mutants are candidate negative feedback regulators, Hsp70ΔFBL – which lacks induction of all four paralogs – shows sustained and elevated HSE-YFP levels following heat shock (Figure 3A, bottom). Thus, induction of Hsp70 is required for HSR deactivation, establishing Hsp70 as a bona fide negative feedback regulator (*17, 18*).

**Figure 3.**
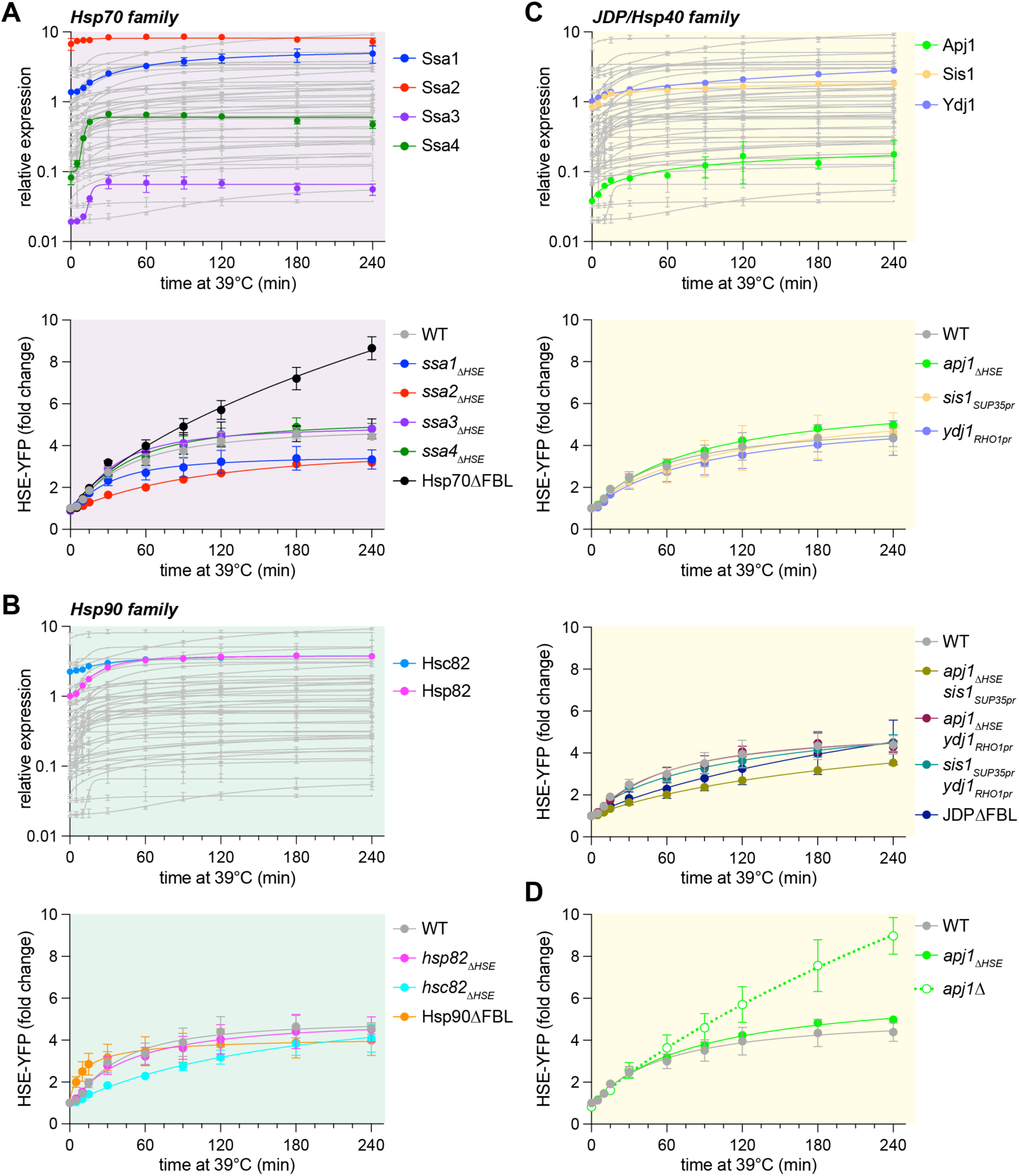
Assessment of functional redundancy among Hsp70, Hsp90, and JDP families. A) Top: Expression of Hsp70 paralogs as measured by P2A-mScarlet over heat shock in color plotted over the full library in gray. Bottom: HSE-YFP fold change for single Hsp70 induction ΔHSE mutants and Hsp70ΔFBL, in which none of the paralogs can be induced over a heat shock time course. Each data point represents the mean and standard deviation of three biological replicates. B) As in (A) but for the Hsp90 paralogs. C) Top as in (A) but for the JDPs. Middle: Individual JDP induction mutants. Bottom: Combined JDP induction mutants. D) HSE-YFP fold change over a heat shock time course for WT, *apj1ΔHSE*, and apj1Δ. Each data point represents the mean and standard deviation of three biological replicates.

Like the Hsp70 paralogs, the two genes encoding Hsp90, *HSC82* and *HSP82*, also differ in their expression dynamics. While the basal expression of Hsc82 > Hsp82, their expression levels converge over the heat shock time course (Figure 3B, top). To disrupt Hsp90 induction without disturbing its basal expression, we fused *HSC82* to the constitutive *TDH3* promoter matching the combined basal level of both paralogs in a strain deleted for the endogenous copies, generating Hsp90ΔFBL. As opposed to Hsp70ΔFBL, Hsp90ΔFBL cells were able to deactivate the HSR upon sustained heat shock (Figure 3B, bottom). As such, Hsp90 does not fulfill the criteria of a negative feedback regulator of the HSR.

Due to our previous result demonstrating that Sis1 is not a feedback regulator or the HSR (*18*), we wondered whether the other cytosolic JDPs encoded in the HSR regulon could be providing feedback. Ydj1 and Sis1 show comparable basal and induced expression levels, while Apj1 is expressed an order of magnitude lower (Figure 3C, top). Previously, we engineered a strain expressing the only copy of *SIS1* from the *SUP35* promoter to disrupt induction upon heat shock while maintaining its high basal levels (*18*). We employed a similar promoter swapping strategy to set expression of Ydj1 near its basal levels in wild type cells by expressing it from the *RHO1* promoter (Figure S3A, B). Since Apj1 is negligibly expressed under basal conditions, the ΔHSE mutant sufficed (Figure S3C, D). All these individual induction mutants showed wild type-like HSE-YFP induction profiles over a heat shock time course (Figure 3C, middle). Next, we disrupted induction of these JDPs in all pairwise combinations and all three at once. Like the single mutants, the double induction mutants – and even the triple mutant termed JDPΔFBL – all induced HSE-YFP over a heat shock time course comparably to wild type (Figure 3C, bottom). These data indicate that induction of cytosolic JDPs is dispensable for feedback regulation of the HSR.

While initially characterizing the strains to perform these experiments, we observed that, in contrast to the *apj1ΔHSE* strain and the triple JDPΔFBL strain, complete deletion of the gene encoding Apj1 had a pronounced effect on HSR dynamics. In *apj1Δ* cells, HSE-YFP levels are modestly elevated under basal conditions and induced and sustained during heat shock at substantially elevated levels (Figure 3D). Thus, while Apj1 is not a negative *feedback* regulator of the HSR, it is a negative regulator; its presence at basal levels is required for deactivation of the HSR. So, while JDPs are not feedback regulators, two different JDPs negatively regulate the HSR: Sis1 under basal conditions and Apj1 during sustained heat shock.

### Growth measurements of HSR induction mutants at elevated temperature

Since only a small fraction of the HSR regulon confers negative feedback on the pathway during heat shock, we hypothesized that the induction of additional targets would be important for fitness at elevated temperature. Notably, we previously found that Hsp70ΔFBL – which has a strong HSR feedback phenotype – grows comparably to wild type at elevated temperature, while induction of Sis1 – which is dispensable for feedback regulation of the HSR – has a fitness defect during the diauxic shift, indicating that feedback and fitness can be uncoupled (*18*). To determine whether ΔHSE mutants have altered growth during heat shock, we measured quantitative growth curves for each mutant in the library relative to wild type cells in control and elevated temperature growth regimes (Figure 4A). Nine induction mutants had reduced maximal growth rates at 37°C relative to wild type, those affecting expression of Ydj1, Apj1, Gre3, Pin3, Ubi4, Ira2, Apa1, Hsp30, and Fes1 (Figure 4B). In addition, these mutants along with induction mutants of Sis1 and the mitochondrial chaperones showed growth phenotypes in the late stage of growth corresponding to the diauxic shift (Figure 4B). Of the nine mutants with reduced log phase growth, five were also candidate negative feedback regulators as defined by the HSE-YFP reporter assay (Figure 4C). Of all the induction mutants, only *hsp42ΔHSE* showed increased Hsf1 activity without a fitness defect, like Hsp70ΔFBL. Thus, additional members of the HSR regulon confer fitness at elevated temperature. However, more than half the HSR induction mutants display neither feedback nor fitness defects.

**Figure 4.**
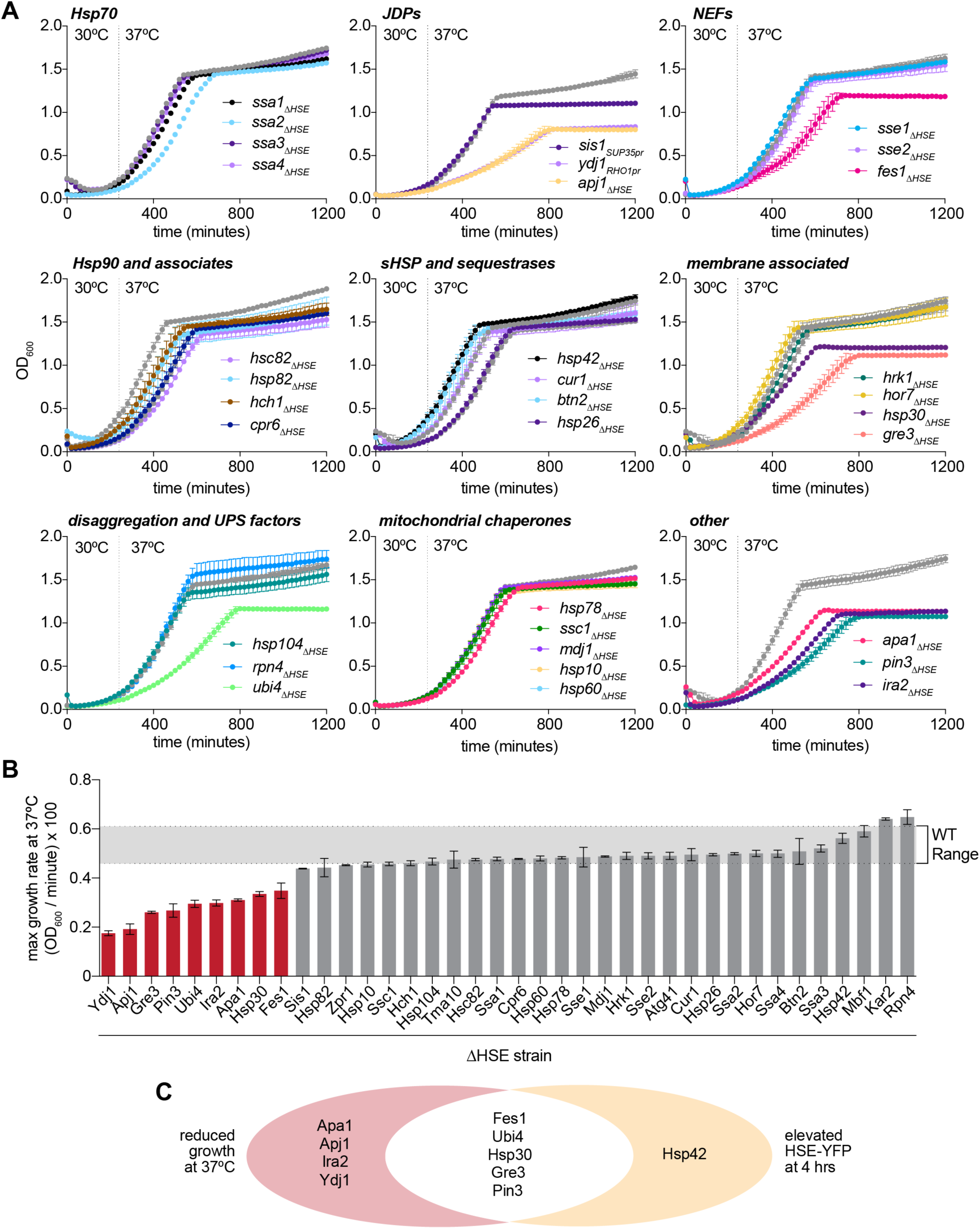
Growth measurements of ΔHSE mutants. A) Quantitative growth assay of Hsf1 target induction mutants grown first at 30°C and switch to 37°C at the designated time, grouped by known cellular function. Each data point represents the mean optical density (OD600) and error bars represent the standard deviation of the two biological replicates. B) The log phase growth rate of each induction mutant (maximum derivative of the OD600 curve). Bar height represents the mean of two biological replicates and error bar represents the range. C) Venn diagram of ΔHSE mutants with growth and HSE-YFP phenotypes.

### Subcellular localization of chaperones in select induction mutants

To determine whether the mutants with both feedback and fitness phenotypes – *fes1ΔHSE*, *ubi4ΔHSE*, *pin3ΔHSE*, and *gre3ΔHSE* – display hallmarks of altered proteostasis, we imaged Hsp104-mKate, which marks cytosolic condensates upon heat shock (*17*). We quantified the fraction of Hsp104-mKate localized to condensates in single cells under basal conditions and following 60 minutes of heat shock, a time point in which the feedback loops have been activated and the cells have largely restored proteostasis. In wild type cells, Hsp104-mKate is diffuse in the cytosol under basal conditions, and by 60 minutes of heat shock, less than 15% of Hsp104 remains condensed in the average cell (Figure 5A). In contrast, all four of the induction mutants with feedback and fitness phenotypes also showed elevated Hsp104 condensation under basal conditions. Three of them – *fes1ΔHSE*, *ubi4ΔHSE*, *pin3ΔHSE* – also displayed large Hsp104-mKate condensates after 60 minutes of heat shock. These imaging data reveal that cytosolic proteostasis is disrupted in these mutants, suggesting that these factors impinge on Hsf1 activity indirectly.

**Figure 5.**
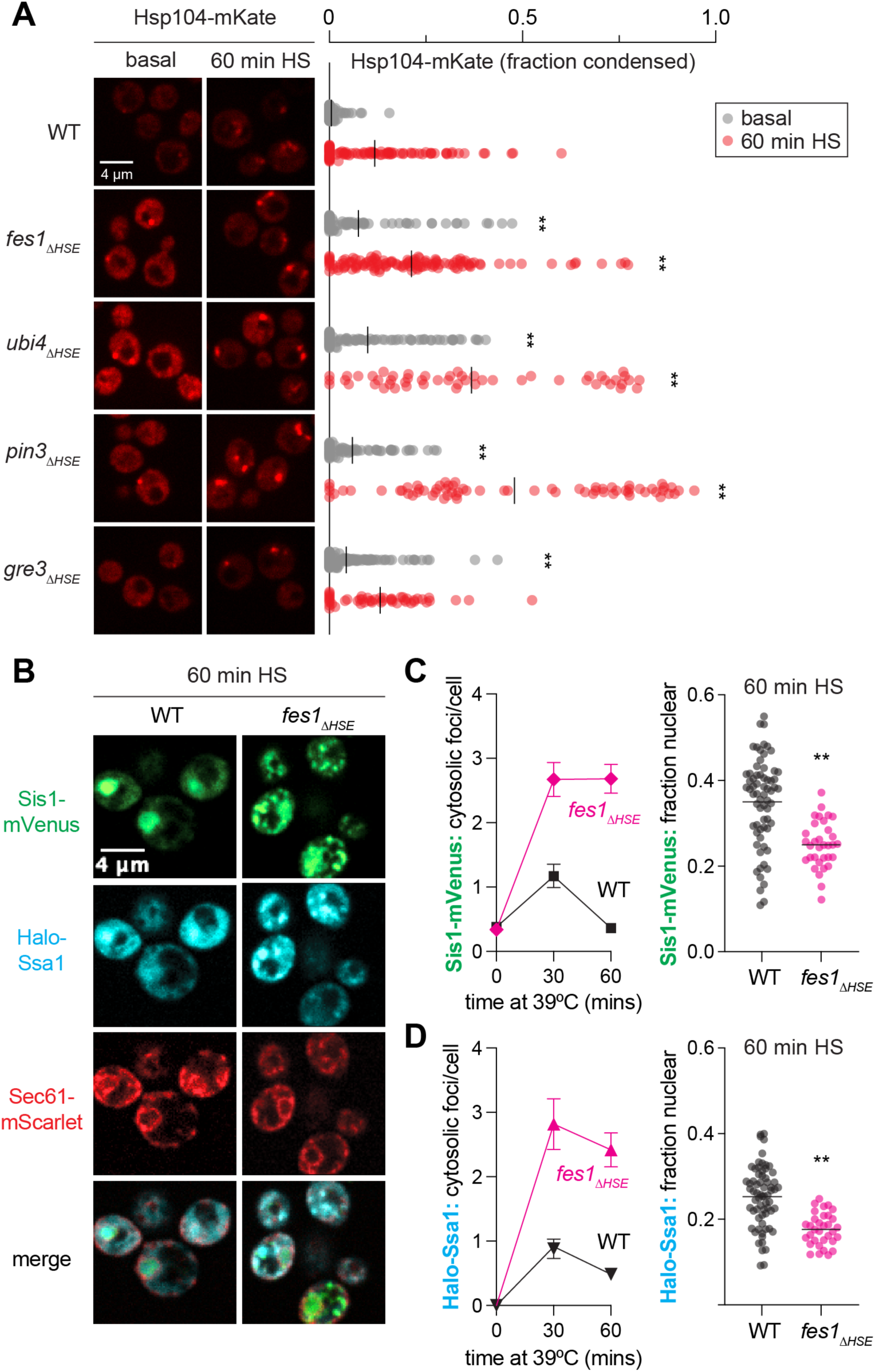
Subcellular localization of chaperones in select induction mutants. **A)** Spinning disk confocal images of wild type and ΔHSE mutants expressing Hsp104-mKate. Scale bar is 4 µm. Fraction of Hsp104-mKate signal condensed in each cell is quantified. Statistics: ** p < 0.01, two-tailed p-value generated from unpaired t-test. **B)** Spinning disk confocal images of induction mutants expressing Halo-Ssa1 to monitor Hsp70 and Sis1-mVenus, and endogenously tagged Sec61-mScarlet to mark the nuclear boundary and cell cortex. Scale bar is 4 um. **C)** Single cell count of condensates and quantification of nuclear localization of Sis1-mVenus during heat shock at 39°C. Each data point represents the average number of foci per cell and standard deviation of those counts for > 25 cells. Statistics: ** p < 0.01, two-tailed p-value generated from unpaired t-test. **D)** As in (C) but for Halo-Ssa1.

Since Hsp104 cooperates with Hsp70 and its co-chaperones to disperse substrates (*32, 45*), we hypothesized that the sustained cytosolic Hsp104 foci we observed would correspond to an increase in cytosolic localization of the key regulators of the HSR – Hsp70 and Sis1. This could result in decreased localization of Hsp70 and Sis1 to the nucleus leading to de-repression of Hsf1 and activation of the HSR. To test this, we generated a *fes1ΔHSE* strain – the induction mutant with the strongest HSE-YFP phenotype – expressing Halo-Ssa1 to image Hsp70, Sis1-mVenus, and Sec61-mScarlet to mark the nuclear boundary and cell cortex. Both Halo-Ssa1 and Sis1-mVenus localized to cytoplasmic condensates upon heat shock in wild type and *fes1ΔHSE* cells, but Ssa1 and Sis1 formed a greater number of condensates in *fes1ΔHSE* cells than in wild type cells and the condensates persisted longer (Fig 5B-D). Correspondingly, *fes1ΔHSE* cells displayed a significantly reduced fraction of Halo-Ssa1 and Sis1-mVenus localized to the nucleus than wild type cells following 60 minutes of heat shock. Thus, consistent with its biochemical function as a nucleotide exchange factor for Hsp70 (*20, 46*), induction of Fes1 is required to release Hsp70 from the cytosolic condensates to deactivate the HSR in the nucleus (Figure 6A).

**Figure 6.**
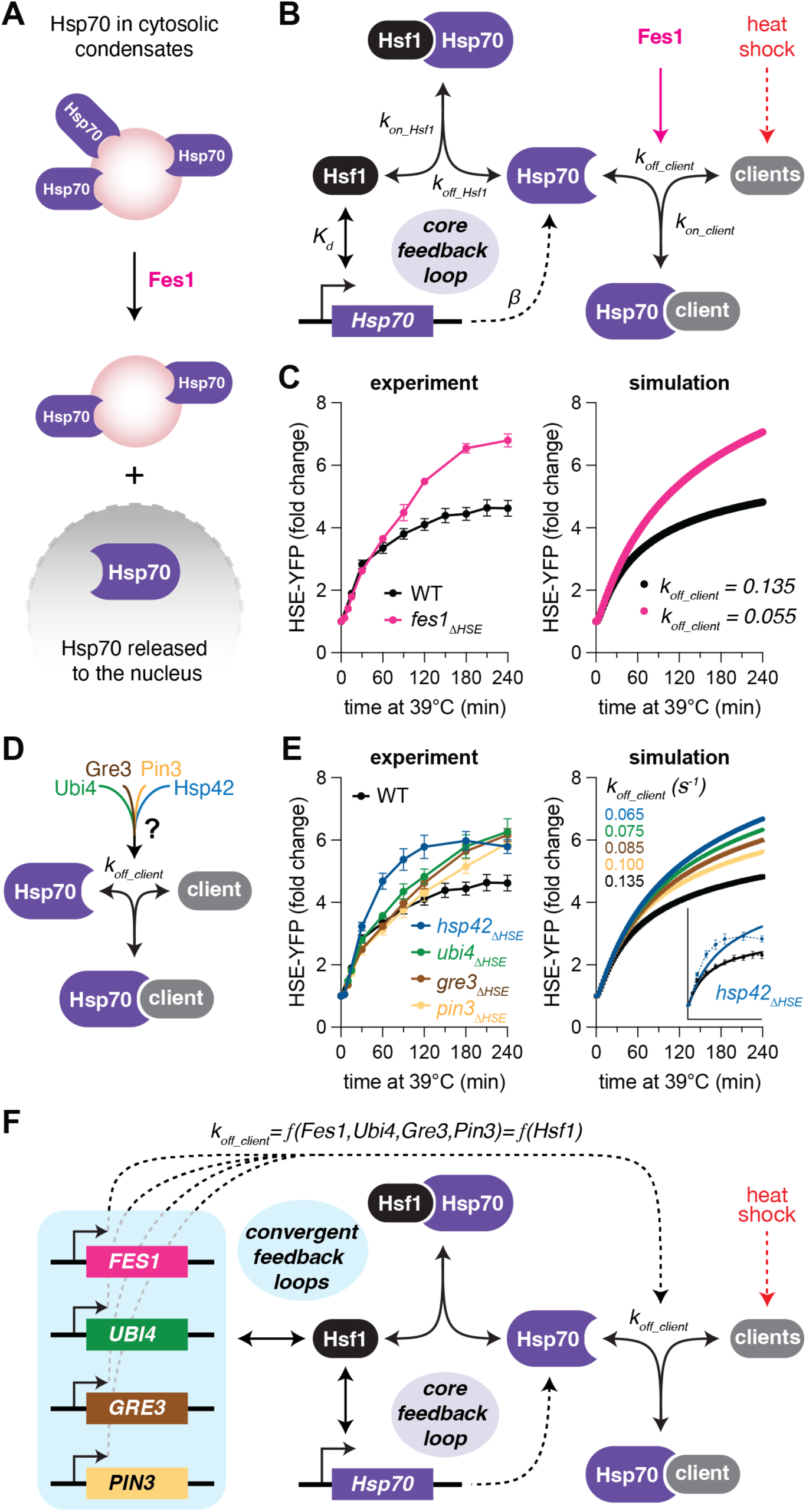
Model of HSR with feedback regulation of Hsp70 expression and localization. A) Schematic of the role of Fes1 in releasing Hsp70 from cytosolic condensates. B) Schematic of mathematical model of the HSR. C) Experimental data and model simulations of HSE-YFP levels over heat shock time courses in wild type and *fes1ΔHSE* cells. Experimental data points and error bars are mean and standard deviation of three biological replicates. D) Schematic of convergence of auxiliary feedback factors on Hsp70 client release. E) As in (C) but for the wild type, *ubi4ΔHSE*, *gre3ΔHSE*, *pin3ΔHSE*, and *hsp42ΔHSE*. F) Schematic of the convergence of the auxiliary feedback factors into a single parameter in the mathematical model.

### Mathematical model with feedback regulation of both Hsp70 expression and localization

We have refined a mathematical model of the HSR over the course of several studies (*17–19, 21*). The model is based on a core two-component feedback loop, in which Hsf1 activates expression of Hsp70 while Hsp70 represses the activity of Hsf1, that controls the dynamics of the transcriptional regulon as measured by the HSE-YFP reporter. Upon heat shock, we simulate a temperature-dependent decrease in the spontaneous folding rate of newly synthesized proteins, resulting in the accumulation of “clients” for Hsp70. Via an affinity switch that captures the titration of the JDP Sis1 away from Hsf1 by accumulated clients, Hsp70 dissociates from Hsf1, and Hsf1 induces expression of more Hsp70 until the system adapts to a new steady state (*21*).

To incorporate the roles of the novel feedback regulators, we first fit the model to *fes1ΔHSE*. Architecturally, we modeled the action of Fes1 as promoting the productive release of Hsp70 from client proteins (Figure 6B). Mathematically, we modeled the *fes1ΔHSE* strain by reducing the value of the parameter describing this productive release rate until we maximized the goodness of fit (Figure S4). Simulation of heat shock time courses in wild type and *fes1ΔHSE* cells quantitatively recapitulated the HSE-YFP induction dynamics we measured experimentally (Figure 6C). Aside from adjusting this single parameter, this updated model of the HSR required no further parameter adjustments nor any structural changes from the previous version. Indeed, tuning this same parameter also enabled us to recapitulate the dynamics of *gre3ΔHSE*, *pin3ΔHSE*, and *ubi4ΔHSE* (Figure 6D, E), supporting the notion that these factors converge with Fes1 to enable efficient restoration of cytosolic proteostasis and Hsp70 client release. Notably, modulating this parameter failed to recapitulate the dynamic HSE-YFP response we observed in the *hsp42ΔHSE* mutant (Figure 6E), which is hyperactive at early time points relative to wild type and the other mutants. Consistent with its lack of growth phenotype, the modeling indicates that Hsp42 impinges on the HSR via a distinct mechanism than the other feedback mutants.

Taken together, these experimental and modeling results suggest that, except Hsp42, the additional feedback regulators converge on the HSR by modulating the nuclear availability of Hsp70. Without any new parameters or new species, we can interpret the mathematic model in the context of the new results: the HSR is governed by a core feedback loop controlling Hsp70 expression supplemented by an auxiliary feedback loop – into which multiple targets converge – controlling Hsp70 client release and thus its subcellular localization (Figure 6F).

### Regulon-wide screen for Hsf1 feedback regulators during ethanol stress

The HSR is activated by a wide range of stressors beyond heat shock, including ethanol, reactive oxygen species, and specific perturbations to the proteostasis network (*1, 43, 47*). To determine whether additional factors may participate in feedback regulation of the HSR under a different condition, we repeated the screen in of the ΔHSE mutants following exposure to ethanol. In wild type cells expressing the HSE-YFP reporter, we observed dose- and time-dependent induction of the HSR as a function of ethanol concentration, with half-maximal activity (EC50) at 6.9% ethanol and corresponding inhibition of growth (IC50) at 6.6% ethanol (Figure 7A, B, Figure S5A, B). We settled on a dose of 8.5% – above the EC50 and used recently to study the HSR (*48*) – to perform the time course screen of the library (Figure 7C).

**Figure 7.**
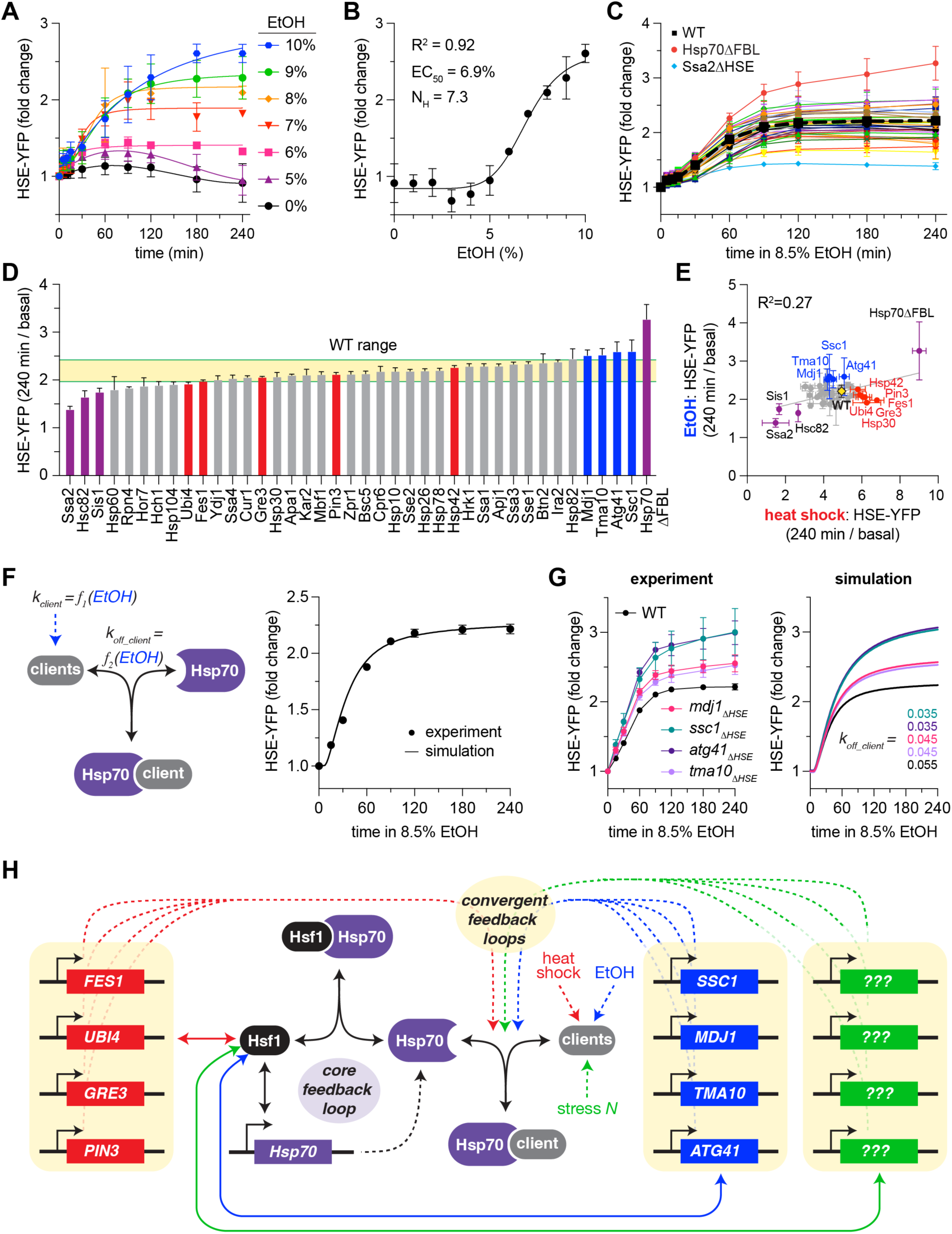
Regulon-wide screen for Hsf1 feedback regulators during ethanol stress. **A)** Time dependent dose response of the HSE-YFP reporter to ethanol (EtOH) in wild type cells. Each data point and error bar represents the mean and standard deviation of three biological replicates. 6-10% ethanol time courses were well-fit by sigmoidal curves, but 5% and 0% had to be fit with polynomial functions. **B)** HSE-YFP levels at 4 hours were fit to a Hill function. **C)** HSE-YFP fold change over a time course of treatment with 8.5% EtOH for all ΔHSE mutants, measured by flow cytometry. Dashed black line represents WT average. Each data point represents the mean and standard deviation of three biological replicates. **D)** HSE-YFP fold change after four hours of 8.5% EtOH for each ΔHSE mutant and the Hsp70ΔFBL mutant. WT range is shown in yellow. Bars show the mean of 3 biological replicates, error bars the standard deviation. Purple bars show outliers common to heat shock and EtOH; blue bars show outliers in EtOH; red bars show outliers in heat shock. **E)** Scatter plot of HSE-YFP fold change after four hours in heat shock versus EtOH for the ΔHSE mutants. **F)** Schematic and simulation of the mathematical model re-parameterized for the HSE-YFP response to ethanol in wild type cells. **G)** Experimental data and model simulations of HSE-YFP levels over EtOH time courses in wild type and auxiliary feedback mutant cells. Experimental data points and error bars are mean and standard deviation of three biological replicates. **H)** Model of how different subsets of the HSR regulon converge to function as the auxiliary feedback to control Hsp70 subcellular localization under different conditions to confer adaptive flexibility.

Like the results in response to heat shock, Hsp70ΔFBL had the highest level of induction after four hours of ethanol exposure while *ssa2ΔHSE, sis1ΔHSE,* and *hsc82ΔHSE* all showed the lowest levels of induction due to their high basal levels (Figure 7D, purple bars). However, the other induction mutants that showed elevated HSE-YFP levels following heat shock were indistinguishable from wild type in response to ethanol (Figure 7D, red bars). Instead, four new mutants – *ssc1ΔHSE*, *mdj1ΔHSE*, *atg41ΔHSE*, and *tma10ΔHSE* – show phenotypes consistent with disrupted feedback (Figure 7D, blue bars). Further supporting a distinct feedback network in response to ethanol and heat shock, we found that across the induction mutant library, the ratio of fold change at four hours following heat shock and ethanol for each mutant varied substantially (R^2^ = 0.27, Figure 7E). These data suggest that the auxiliary feedback loop may be comprised of condition-specific subsets of the HSR regulon.

Despite the distinct subsets of feedback regulators following heat and ethanol, we hypothesized that we could nonetheless recapitulate the altered dynamics of the ethanol-specific feedback candidates by modeling them as converging on the auxiliary feedback loop controlling Hsp70 client release. To this end, we first reconfigured the model to simulate ethanol exposure rather than heat shock. Without altering the architecture of the model or the parameters describing the core feedback loop, we accomplished this by adjusting the parameters describing the rate of client production and the interaction of clients with Hsp70 until the model could faithfully reproduce the HSE-YFP dynamics of wild type cells (Figure 7F). Indeed, with this model of the HSR induced by ethanol, we were able to account for the altered HSE-YFP dynamics of the ethanol-specific feedback candidates by adjusting the single parameter describing Hsp70 client release (Figure 7G). Given the broad range of environmental conditions that activate the HSR, additional perturbations may likewise require induction of distinct subsets of the HSR regulon to productively engage different clients and promote Hsp70 release (Figure 7H).

## DISCUSSION

In this study, we comprehensively dissected transcriptional feedback in the HSR. While negative feedback loops have long been appreciated to architecturally organize cellular stress response pathways, we present here the first regulon-wide screen of response dynamics to identify feedback regulators in an environmental stress pathway. The major conclusion we draw is that the HSR is governed by a core negative feedback loop that determines Hsp70 expression augmented by a condition-specific auxiliary feedback loop that controls Hsp70 subcellular localization. In cell biological terms, the results support a model in which the availability of Hsp70 in the nucleoplasm largely determines the transcriptional output of the HSR across conditions and timescales. We anticipate that this architecture, in which the availability of a key pathway regulator is controlled via expression-level feedback and fine-tuned by spatial feedback, will characterize adaptive responses beyond the HSR.

Before screening the HSR regulon for feedback factors, we first characterized the induction dynamics of each of the 42 genes in the regulon. Analysis of the resulting dataset revealed that the HSR genes span a linear spectrum ranging from rapid/transient induction to slow/sustained induction (Figure 1G). Thus, while the transcription factor Hsf1 is evidently capable of regulating different genes across an expression range spanning four orders of magnitude, the one-dimensional manifold of the gene induction profiles suggests that simple underlying constraints determine the variation in expression across the regulon. Our previous finding that the level of Hsf1 binding at each gene can be predicted by the affinity of the HSE in the promoter for Hsf1 and the chromatin accessibility of the locus may provide the mechanistic explanation (*5*). In the context of HSR feedback regulation, it is notable that the *SSA2/3/4* paralogs of Hsp70 – components of the core negative feedback – are among the genes nearest the rapid/transient end of the induction spectrum, while *FES1* and the JDPs *SIS1*, *APJ1*, *YDJ1* – the strongest auxiliary feedback regulator and non-feedback regulators, respectively – are among the genes nearest the slow/sustained end (Figure 1G).

In addition to identifying Fes1, the ΔHSE mutant screen for altered HSR dynamics following heat shock revealed several other negative feedback regulators: Ubi4, Gre3, Pin3, and Hsp42. Except *hsp42ΔHSE*, the feedback mutants exhibited reduced growth rates and disrupted cytosolic proteostasis (Figure 4C, Figure 5A). Moreover, their HSE-YFP dynamics could be recapitulated by altering the value of a single parameter describing the rate of client release by Hsp70 in a mathematical model of the HSR (Figure 6F). For *fes1ΔHSE*, we directly demonstrated that Hsp70 remains localized in cytosolic condensates, supporting the modeling results. Since Fes1 functions as a NEF for Hsp70, and nucleotide exchange is coupled to Hsp70 client release (*20*), it is intuitive why Fes1 induction during heat shock would be required for efficient liberation of Hsp70 from cytosolic clients and subsequent nuclear localization. Likewise, induction of *UBI4*, which encodes concatameric ubiquitin (*49, 50*), can be rationalized as important for restoring cytosolic proteostasis due to its central role in the ubiquitin-proteasome system, thereby indirectly impinging on the availability of Hsp70.

The functions of Gre3 and Pin3 in the proteostasis network are less well understood. Gre3 functions as an aldose reductase that acts to convert methylglyoxal generated by glycolysis during stress to pyruvate (*51*). Increased levels of methylglyoxal in *gre3ΔHSE* cells may react with and damage cytosolic proteins, thereby sequestering Hsp70. Pin3 regulates actin nucleation and has been implicated in prion formation (*52, 53*). Perhaps induction of Pin3 is important for actin-dependent adaptive remodeling of cytosolic condensates that somehow serves to free Hsp70. While the details of the mechanisms remain to be resolved, it is likely that these additional feedback regulators are performing independent functions that converge to determine the availability of Hsp70 in the nucleus following heat shock.

The role of induction of Hsp42 in HSR regulation is more mysterious. Unlike the auxiliary feedback mutants – but like the core feedback mutant Hsp70ΔFBL – *hsp42ΔHSE* has no growth phenotype at elevated temperature. Also, relative to the auxiliary feedback mutants, lack of induction of Hsp42 alters the dynamics of the HSE-YFP reporter at earlier time points, and the resulting time course data cannot be fit by adjusting the Hsp70 client release parameter (Figure 6E). Based on our results, we cannot rule out that Hsp42 acts to directly regulate Hsf1, though we have no evidence to support this.

Intriguingly, our screen for feedback mutants during ethanol stress revealed four distinct negative feedback regulators of the HSR: Ssc1, Mdj1, Atg41 and Tma10. Remarkably, these factors are all implicated in regulating mitochondrial homeostasis. Ssc1 and Mdj1 encode a mitochondrially-targeted Hsp70 and JDP, respectively (*54, 55*); Atg41 localizes to the mitochondrial surface and is required for mitophagy (*56*); and while Tma10 has no known function, its paralog Stf2 is known to bind and regulate the F1FO mitochondrial ATP synthase (*57, 58*). Thus, the subset of the HSR regulon that makes up the ethanol-specific auxiliary feedback loop is comprised of genes that reflect the physiological nature of ethanol stress— namely that ethanol is a non-fermentable carbon source metabolized in the mitochondria. These results imply that without induction of mitochondrial-specific factors encoded in the larger HSR regulon, the increased stress the mitochondria experiences in the presence of ethanol spills into the cytosol, titrating Hsp70 away from the nucleus. Consistent with this notion, the mitochondrial unfolded protein response was recently demonstrated to be mediated via titration of cytosolic Hsp70 and subsequent de-repression of Hsf1 in the nucleus (*59*).

Taken all together, we propose a model in which the HSR is dynamically regulated by a core feedback loop driven by induction of Hsp70 expression that acts regardless of the nature of the stress augmented by an auxiliary feedback loop that is condition-specific. In this view, each condition could produce a unique set of clients that would require a distinct subset of the HSR regulon to manage. These condition-specific feedback factors would then converge to enable efficient Hsp70 release back into the free cytosolic pool that can diffuse into the nucleus and repress Hsf1.

What advantages would such a two-tiered feedback architecture comprised of core and auxiliary loops confer to an adaptive response? On evolutionary timescales, this structure allows a simple, invariant network to remain fixed within a population – in this case the Hsp70-Hsf1 negative feedback loop – while providing plasticity such that a secondary, peripheral genetic network tailored to cope with specific environmental fluctuations in any given ecological niche can evolve to fine-tune the output of the core network. In this case, the existence of two distinct subsets of condition-specific HSR effectors reflects an evolutionary past of fluctuating temperatures and carbon sources. The theoretical alternative to this mode of adaptation is a direct rewiring of the core Hsp70-Hsf1 feedback loop in new environments. This would likely render the system susceptible to mutations that come with fitness costs in the presence of new environmental challenges. I.e., it may be a less evolvable mode of adaptation.

Extrapolating from the two conditions we tested here, we suspect that the HSR target genes not implicated in feedback during heat shock or ethanol stress may similarly converge to fine tune Hsp70 subcellular localization in other environmental conditions experienced in the evolutionary history of budding yeast, such as nutrient, pH, and redox fluctuations. The two-tiered feedback architecture of the HSR allows for adaptive flexibility while maintaining a conserved core that was likely already present in – and may have been essential for the evolvability of – the last eukaryotic common ancestor.

## METHODS

### Strain Construction

To create the library of P2A-mscarlet tagged strains, we took advantage of yeast homologous recombination by introducing a P2A-mscarlet-KAN cassette with homologous flanking ends and plated on kanamycin selective media. Successful transformants were verified by PCR and flow cytometry. To create ΔHSE induction mutants, we deleted the 9-25 bp Hsf1-binding consensus sequence by scarless CRISPR-Cas9 targeted deletion (*60*). The closest match to the Hsf1 binding motif (nnTTCnnGAA) was located under the 5 min heat shock Hsf1 ChIP-seq peak from our previous study. Generally, we found one strong consensus sequence under the singular Hsf1 ChIP-peak ahead 200-300 bp ahead of the TSS. We delete the ChIP-verified consensus sequence, in the established library of P2A-mscarlet parent lines. Cell line construction involved cloning a guide RNA, which will target the Hsf1 binding site, into an episomal, URA3-marked, Cas9-containing plasmid. The guide RNA plasmid was co-transformed with a 100 bp double-stranded repair template to repair the double stranded break by homologous recombination, and yeast were plated on ura-selective media. HSE deletion was confirmed by Sanger sequencing. Finally, the cell lines were plated on 5-FOA selective media to expel the Cas9::ura3 plasmid.

Though our P2A-mscarlet reporter strategy was tagless, it caused abnormal basal Hsf1 activity in a few lines, probably because the C terminal linker inhibited protein function. So, we created another library of ΔHSE induction mutants in our original parent line w303a; HSE-mVenus. The library of induction mutants (without the mScarlet reporter) was used in all experiments beyond the initial target gene transcription dynamics study.

### Heat shock and ethanol time courses

Cells were serially diluted and grown overnight on the benchtop at room temperature (25°C) in 1xSDC media. In the morning, cells were transferred to microcentrifuge tubes and aerated by shaking (1250 RPM) at 30°C for 1 hour. Centrifuge tubes were then transferred to a shaking incubator at heat shock temperature --39°C or the indicated concentration of ethanol was added. At each time point, 50 uL of cells were transferred to a 96 well plate of 1xSDC at 50 ug/mL final concentration cycloheximide on the benchtop. After the time course, the plate was incubated at 30°C for 1 hour to promote fluorescent reporter maturation before flow cytometry.

### Flow Cytometry

Cells were measured with the BD Fortessa HTS 4-15 benchtop analyzer at the University of Chicago Cytometry and Antibody Technology Facility. Analysis was completed in FlowJo: fluorescence excitation value for each cell was normalized by side scatter for each cell to filter out signal from dead cells, and the median normalized fluorescence excitation value was calculated for each sample.

### Quantitative Growth Assay

Cells were grown overnight shaking in 1xYPD at 30°C. In the morning, cells were diluted to OD600=0.1 in 1xYPD and transferred to a 48-well plate. Cells were grown while shaking in the SPECTROstar Nano® Absorbance Plate Reader at 30°C for four hours, then at 37°C for 20 hours, and OD600 was measured every 20 minutes. The initial four hour incubation at 30°C before heat shock yields more consistent results across biological replicates. OD600 of the liquid cell culture was measured every twenty minutes over the 24-hour growth assay. Two biological replicates were measured per cell line.

### Heat Shock Time Course and Imaging

Cells were grown in 2xSDC media at 30°C shaking overnight. In the morning, cells were diluted to OD600=0.1 and grown at 30°C shaking for 4 hours to reach log phase growth. 200 uL of cells were transferred to a microcentrifuge tube in the cell shaker at 39°C. At each time point, cells were fixed in 1% paraformaldehyde, then washed and imaged in KPIS media (1.2 M sorbitol, 0.1 M potassium phosphate, pH 7.5). Fixed cells were imaged on the Marianas Leica II Spinning Disk Confocal microscope at the University of Chicago Imaging Core. A single z stack was captured and analyzed for each frame. A minimum of 20 cells were quantified at each time point in ImageJ.

### Image quantification

ImageJ was used for all image quantification. Hsp104 foci in each cell were identified using by Intermodes thresholding of each cell. Hsp70 and Sis1 foci were identified using the Triangle threshold. To determine the bounds of the nucleus, a ROI was drawn by hand based on the bounds of the nuclear membrane, marked by Sec61-mscarlet.

### Mathematical modeling

Modeling was performed as described previously using the same equations (*18*). Best fit parameters were determined by minimizing residual sum squared. All updated parameters are described in the text.

## ACKNOWLEDGEMENTS

We thank Kabir Husain and Arvind Murugan for helpful discussions and unfettered access to their plate readers for the numerous growth curve measurements in this study. We thank Asif Ali for technical advice with imaging and members of the Pincus lab for insightful discussions. We would like to acknowledge the University of Chicago Cytometry and Antibody Technology facility (Facility RRID: SCR_017760) for the substantial usage of the high throughput flow cytometers to enable this project. We would also like to acknowledge the University of Chicago Integrated Light Microscopy Core (RRID: SCR_019197). This work was supported by NIH R01 GM138689 and NSF QLCI QuBBE grant OMA-2121044.

---

**Figure S1.**
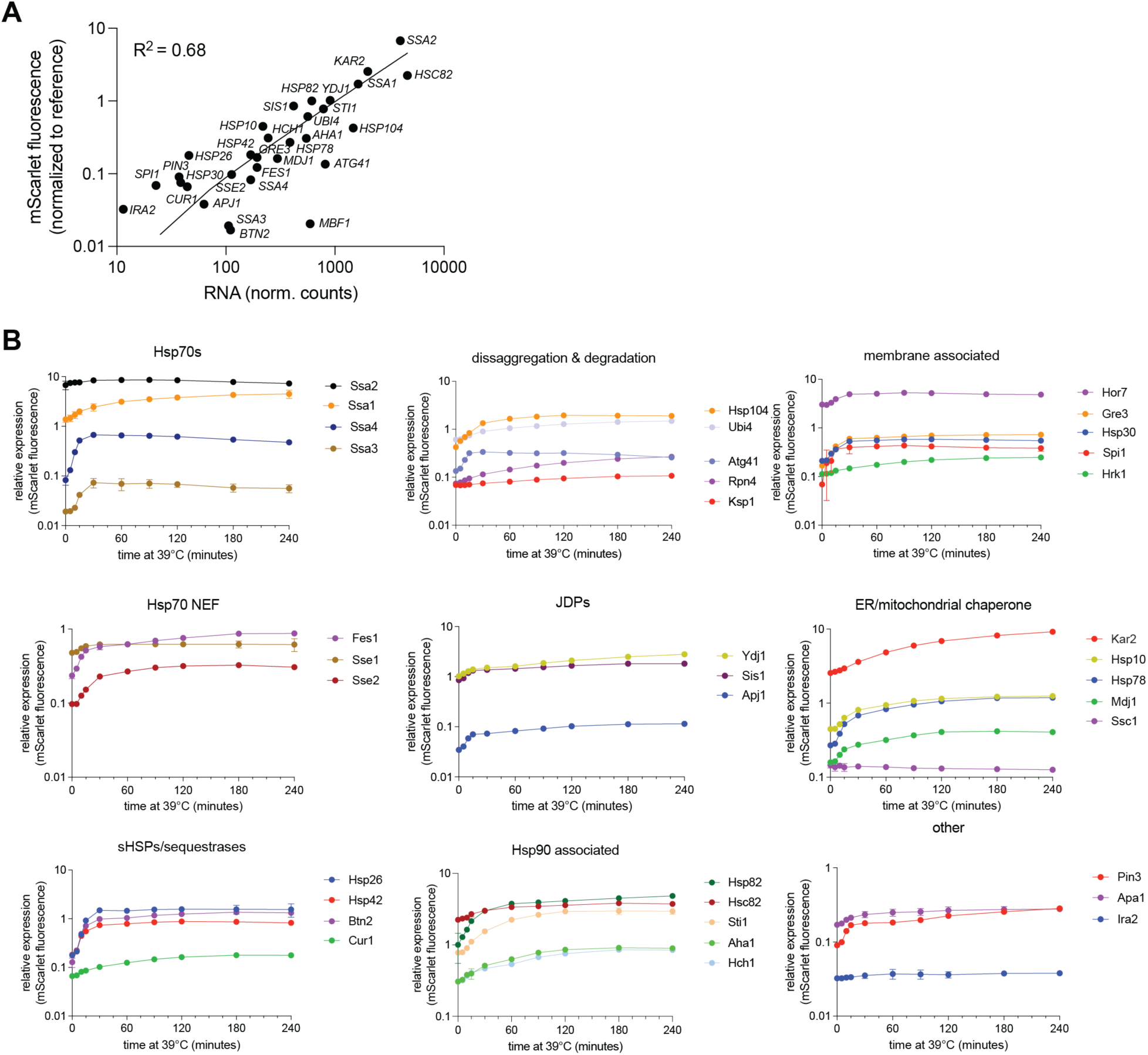
HS-induced expression dynamics of the Hsf1 target genes. **A)** Scatter plot of the expression of the HSR genes as measured by RNA sequencing versus mScarlet expression. **B)** Relative expression dynamics for Hsf1 target genes, captured with mscarlet fluorescent reporter, over a heat shock time course. Hsf1 target genes are grouped by putative gene function. Each data point and error bar represents the mean and standard deviation of 3 biological replicates.

**Figure S2.**
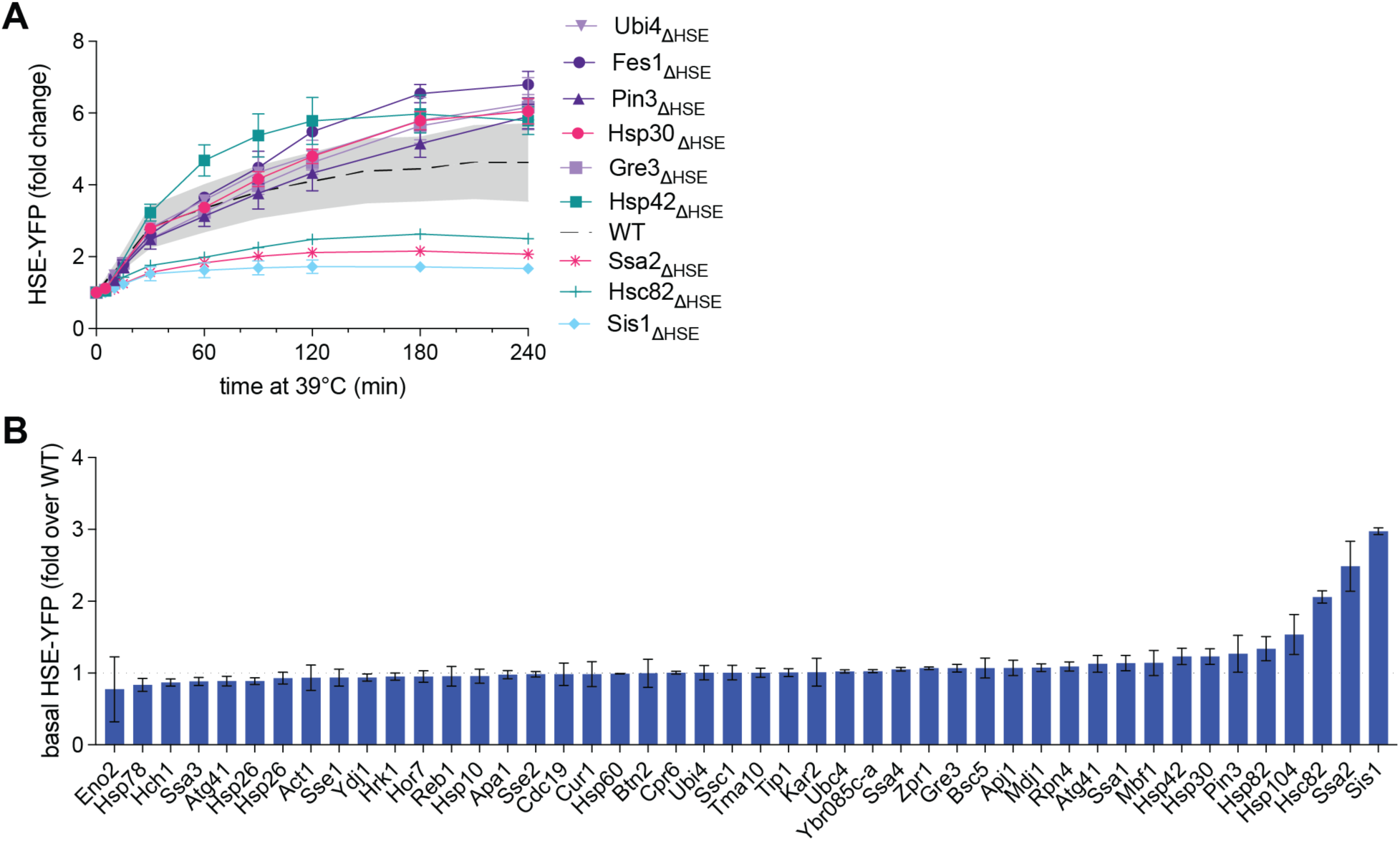
Hsf1 activity dynamics after HSE deletion. **A)** Heat shock time courses for the eight ΔHSE mutants with four-hour HSE-YFP fold change outside of the statistically significant WT range. Each data point represents the mean and standard deviation of three biological replicates. Dotted line represents the average of 45 WT biological replicates, the gray shaded area represents the standard deviation of those replicates. **B)** Relative non-stress levels of HSE-YFP reporter (normalized to WT) in each ΔHSE induction mutant. Each bar height represents the mean of three biological replicates, error bars represent the SD.

**Figure S3.**
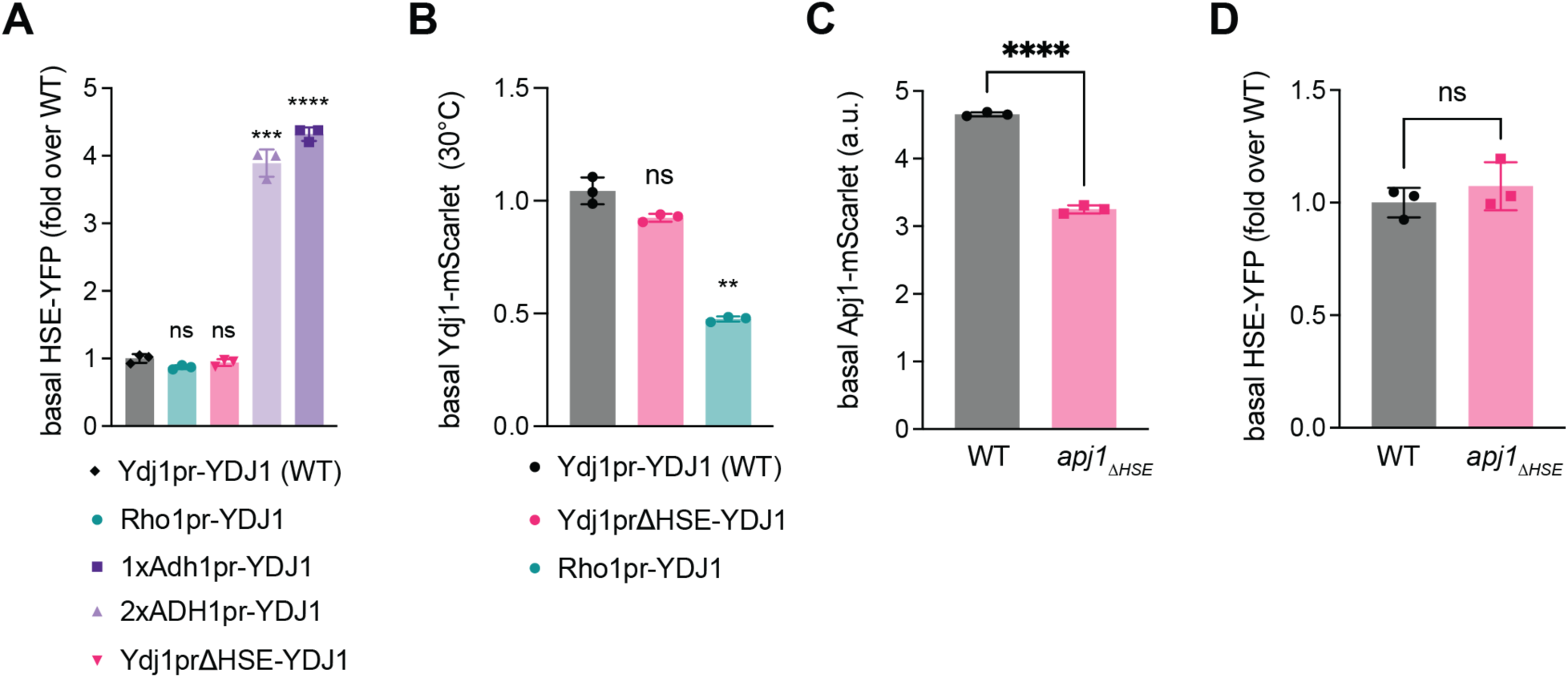
Validating Ydj1 and Apj1 induction mutants. A) Relative basal HSE-YFP (normalized to basal HSE-YFP in WT) when Ydj1 is expressed under non-inducible promoters of various strengths. B) Relative basal Ydj1 expression measured by mScarlet fluorescence. C) Basal Apj1 expression measured by mScarlet fluorescence in *apj1ΔHSE* vs WT. Statistics: p < 0.01. Bar height represents the mean of 3 biological replicates, error bars represent the standard deviation. D) Basal Hsf1 activity measured by HSE-YFP fluorescent reporter in Apj1ΔHSE vs WT. Bar height represents the mean of 3 biological replicates, error bars represent the standard deviation. Statistics: ns is defined as p > 0.05.

**Figure S4.**
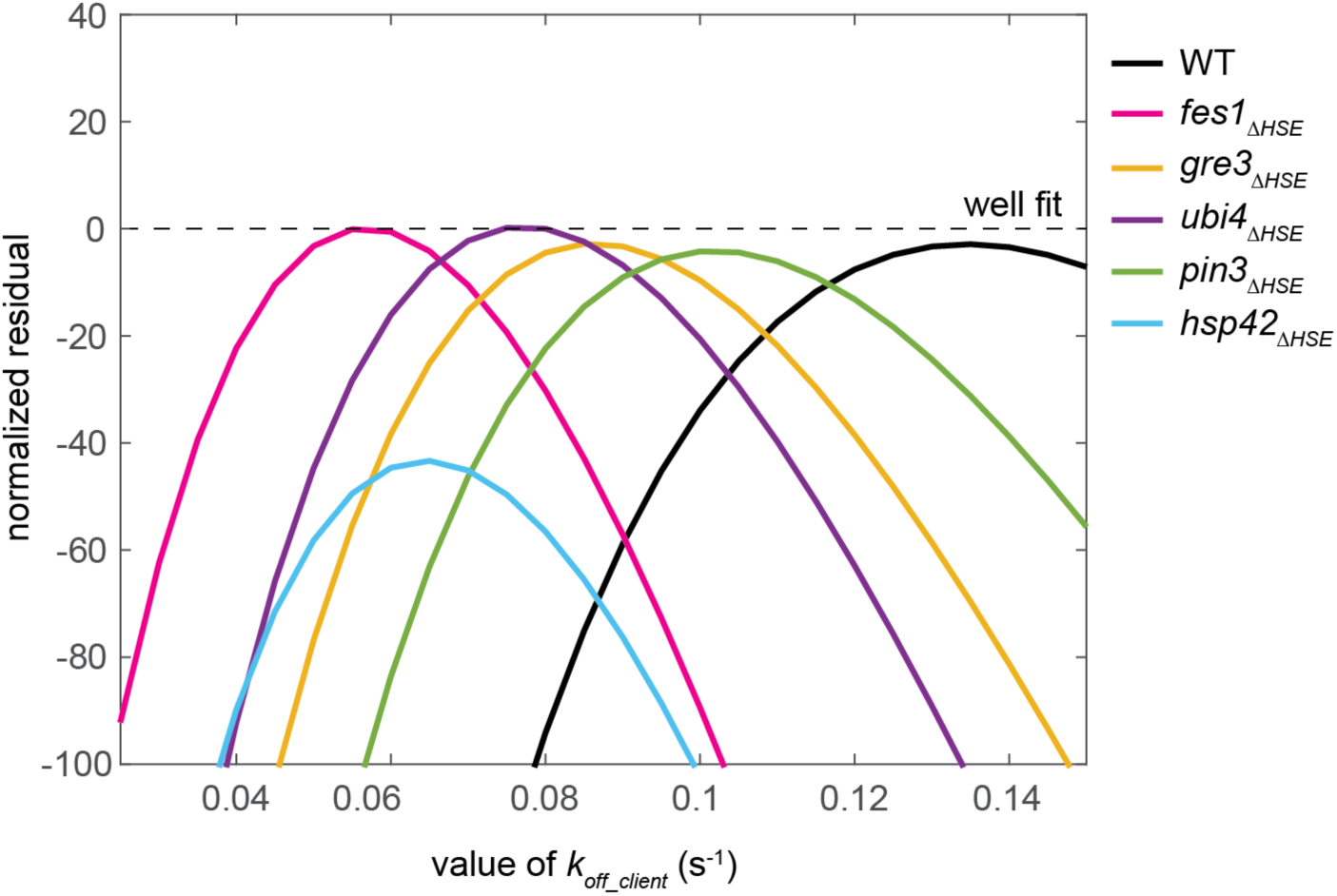
Residuals of model fits. The residual values (quantity left unfit by the model) as a function of the parameter sweeps for each ΔHSE mutant and wild type.

**Figure S5.**
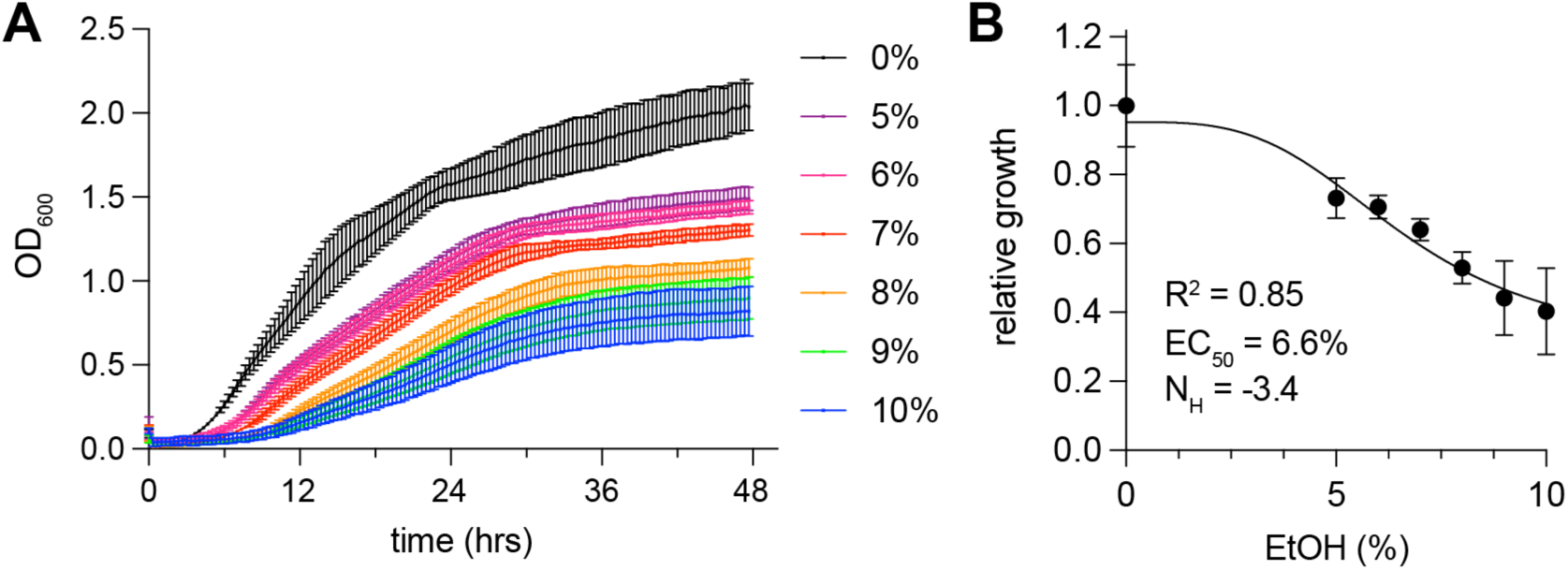
Growth curves of cells in ethanol. **A)** Cells were grown at the indicated concentrations. **B)** Max growth rates were fit to a Hill function.

## REFERENCES

1. D. Pincus, Regulation of Hsf1 and the Heat Shock Response. Adv Exp Med Biol 1243, 41–50 (2020).

2. J. Joutsen, L. Sistonen, Tailoring of Proteostasis Networks with Heat Shock Factors. Cold Spring Harb Perspect Biol 11, (2019).

3. A. S. Kainth, S. Chowdhary, D. Pincus, D. S. Gross, Primordial super-enhancers: heat shock-induced chromatin organization in yeast. Trends Cell Biol 31, 801–813 (2021).

4. E. J. Solis et al., Defining the Essential Function of Yeast Hsf1 Reveals a Compact Transcriptional Program for Maintaining Eukaryotic Proteostasis. Mol Cell 63, 60–71 (2016).

5. D. Pincus et al., Genetic and epigenetic determinants establish a continuum of Hsf1 occupancy and activity across the yeast genome. Mol Biol Cell 29, 3168–3182 (2018).

6. 6. M. J. Alasady, M. L. Mendillo, in HSF1 and Molecular Chaperones in Biology and Cancer, M. L. Mendillo, D. Pincus, R. Scherz-Shouval, Eds. (Springer International Publishing, Cham, 2020), pp. 69–85.

7. M. L. Mendillo et al., HSF1 drives a transcriptional program distinct from heat shock to support highly malignant human cancers. Cell 150, 549–562 (2012).

8. S. Santagata et al., High levels of nuclear heat-shock factor 1 (HSF1) are associated with poor prognosis in breast cancer. Proc Natl Acad Sci U S A 108, 18378–18383 (2011).

9. S. Santagata et al., Tight coordination of protein translation and HSF1 activation supports the anabolic malignant state. Science 341, 1238303 (2013).

10. R. Scherz-Shouval et al., The reprogramming of tumor stroma by HSF1 is a potent enabler of malignancy. Cell 158, 564–578 (2014).

11. C. Dai, L. Whitesell, A. B. Rogers, S. Lindquist, Heat shock factor 1 is a powerful multifaceted modifier of carcinogenesis. Cell 130, 1005–1018 (2007).

12. A. J. Sala, L. C. Bott, R. I. Morimoto, Shaping proteostasis at the cellular, tissue, and organismal level. J Cell Biol 216, 1231–1241 (2017).

13. W. E. Balch, R. I. Morimoto, A. Dillin, J. W. Kelly, Adapting proteostasis for disease intervention. Science 319, 916–919 (2008).

14. D. W. Neef, A. M. Jaeger, D. J. Thiele, Heat shock transcription factor 1 as a therapeutic target in neurodegenerative diseases. Nat Rev Drug Discov 10, 930–944 (2011).

15. R. Gomez-Pastor et al., Abnormal degradation of the neuronal stress-protective transcription factor HSF1 in Huntington’s disease. Nat Commun 8, 14405 (2017).

16. R. Gomez-Pastor, E. T. Burchfiel, D. J. Thiele, Regulation of heat shock transcription factors and their roles in physiology and disease. Nat Rev Mol Cell Biol 19, 4–19 (2018).

17. J. Krakowiak et al., Hsf1 and Hsp70 constitute a two-component feedback loop that regulates the yeast heat shock response. Elife 7, (2018).

18. R. Garde, A. Singh, A. Ali, D. Pincus, Transcriptional regulation of Sis1 promotes fitness but not feedback in the heat shock response. Elife 12, (2023).

19. X. Zheng et al., Dynamic control of Hsf1 during heat shock by a chaperone switch and phosphorylation. Elife 5, (2016).

20. A. E. Masser et al., Cytoplasmic protein misfolding titrates Hsp70 to activate nuclear Hsf1. Elife 8, (2019).

21. Z. A. Feder et al., Subcellular localization of the J-protein Sis1 regulates the heat shock response. J Cell Biol 220, (2021).

22. A. Ali et al., Adaptive preservation of orphan ribosomal proteins in chaperone-dispersed condensates. Nature Cell Biology 25, 1691–1703 (2023).

23. S. Chowdhary, A. S. Kainth, D. Pincus, D. S. Gross, Heat Shock Factor 1 Drives Intergenic Association of Its Target Gene Loci upon Heat Shock. Cell Rep 26, 18–28 e15 (2019).

24. S. Chowdhary, A. S. Kainth, S. Paracha, D. S. Gross, D. Pincus, Inducible transcriptional condensates drive 3D genome reorganization in the heat shock response. Mol Cell 82, 4386–4399 e4387 (2022).

25. S. W. Kmiecik, L. Le Breton, M. P. Mayer, Feedback regulation of heat shock factor 1 (Hsf1) activity by Hsp70-mediated trimer unzipping and dissociation from DNA. EMBO J 39, e104096 (2020).

26. S. W. Kmiecik, M. P. Mayer, Molecular mechanisms of heat shock factor 1 regulation. Trends Biochem Sci 47, 218–234 (2022).

27. H. Zhang et al., Reversible phase separation of HSF1 is required for an acute transcriptional response during heat shock. Nature Cell Biology 24, 340–352 (2022).

28. D. Tauber et al., Modulation of RNA Condensation by the DEAD-Box Protein eIF4A. Cell 180, 411–426 e416 (2020).

29. P. Yang et al., G3BP1 Is a Tunable Switch that Triggers Phase Separation to Assemble Stress Granules. Cell 181, 325–345 e328 (2020).

30. H. Glauninger, C. J. Wong Hickernell, J. A. M. Bard, D. A. Drummond, Stressful steps: Progress and challenges in understanding stress-induced mRNA condensation and accumulation in stress granules. Mol Cell 82, 2544–2556 (2022).

31. V. Cherkasov et al., Coordination of translational control and protein homeostasis during severe heat stress. Curr Biol 23, 2452–2462 (2013).

32. H. Yoo, J. A. M. Bard, E. V. Pilipenko, D. A. Drummond, Chaperones directly and efficiently disperse stress-triggered biomolecular condensates. Mol Cell 82, 741–755 e711 (2022).

33. D. Mateju et al., An aberrant phase transition of stress granules triggered by misfolded protein and prevented by chaperone function. EMBO J 36, 1669–1687 (2017).

34. J. A. Kolhe, N. L. Babu, B. C. Freeman, The Hsp90 molecular chaperone governs client proteins by targeting intrinsically disordered regions. Mol Cell 83, 2035–2044 e2037 (2023).

35. S. B. Miller et al., Compartment-specific aggregases direct distinct nuclear and cytoplasmic aggregate deposition. EMBO J 34, 778–797 (2015).

36. S. B. Miller, A. Mogk, B. Bukau, Spatially organized aggregation of misfolded proteins as cellular stress defense strategy. J Mol Biol 427, 1564–1574 (2015).

37. S. Escusa-Toret, W. I. Vonk, J. Frydman, Spatial sequestration of misfolded proteins by a dynamic chaperone pathway enhances cellular fitness during stress. Nat Cell Biol 15, 1231–1243 (2013).

38. E. M. Sontag, R. S. Samant, J. Frydman, Mechanisms and Functions of Spatial Protein Quality Control. Annu Rev Biochem 86, 97–122 (2017).

39. E. M. Sontag et al., Nuclear and cytoplasmic spatial protein quality control is coordinated by nuclear-vacuolar junctions and perinuclear ESCRT. Nat Cell Biol 25, 699–713 (2023).

40. R. Babazadeh et al., Syntaxin 5 Is Required for the Formation and Clearance of Protein Inclusions during Proteostatic Stress. Cell Rep 28, 2096–2110 e2098 (2019).

41. W. J. de Jonge et al., Molecular mechanisms that distinguish TFIID housekeeping from regulatable SAGA promoters. EMBO J 36, 274–290 (2017).

42. X. Zheng et al., Hsf1 Phosphorylation Generates Cell-to-Cell Variation in Hsp90 Levels and Promotes Phenotypic Plasticity. Cell Rep 22, 3099–3106 (2018).

43. B. D. Alford et al., ReporterSeq reveals genome-wide dynamic modulators of the heat shock response across diverse stressors. Elife 10, (2021).

44. O. Brandman et al., A ribosome-bound quality control complex triggers degradation of nascent peptides and signals translation stress. Cell 151, 1042–1054 (2012).

45. K. A. Tipton, K. J. Verges, J. S. Weissman, In vivo monitoring of the prion replication cycle reveals a critical role for Sis1 in delivering substrates to Hsp104. Mol Cell 32, 584–591 (2008).

46. N. K. Gowda et al., Cytosolic splice isoform of Hsp70 nucleotide exchange factor Fes1 is required for the degradation of misfolded proteins in yeast. Mol Biol Cell 27, 1210–1219 (2016).

47. B. D. Alford, O. Brandman, Quantification of Hsp90 availability reveals differential coupling to the heat shock response. J Cell Biol 217, 3809–3816 (2018).

48. L. S. Rubio, S. Mohajan, D. S. Gross, Ethanol stress induces transient restructuring of the yeast genome yet stable formation of Hsf1 transcriptional condensates. bioRxiv, 2023.2009.2028.560064 (2023).

49. D. Finley, E. Ozkaynak, A. Varshavsky, The yeast polyubiquitin gene is essential for resistance to high temperatures, starvation, and other stresses. Cell 48, 1035–1046 (1987).

50. E. Ozkaynak, D. Finley, M. J. Solomon, A. Varshavsky, The yeast ubiquitin genes: a family of natural gene fusions. EMBO J 6, 1429–1439 (1987).

51. J. Aguilera, J. A. Prieto, The Saccharomyces cerevisiae aldose reductase is implied in the metabolism of methylglyoxal in response to stress conditions. Curr Genet 39, 273–283 (2001).

52. T. A. Chernova et al., Prion induction by the short-lived, stress-induced protein Lsb2 is regulated by ubiquitination and association with the actin cytoskeleton. Mol Cell 43, 242–252 (2011).

53. A. Madania et al., The Saccharomyces cerevisiae homologue of human Wiskott-Aldrich syndrome protein Las17p interacts with the Arp2/3 complex. Mol Biol Cell 10, 3521–3538 (1999).

54. N. Rowley et al., Mdj1p, a novel chaperone of the DnaJ family, is involved in mitochondrial biogenesis and protein folding. Cell 77, 249–259 (1994).

55. E. A. Craig, J. Kramer, J. Kosic-Smithers, SSC1, a member of the 70-kDa heat shock protein multigene family of Saccharomyces cerevisiae, is essential for growth. Proc Natl Acad Sci U S A 84, 4156–4160 (1987).

56. Z. Yao, E. Delorme-Axford, S. K. Backues, D. J. Klionsky, Atg41/Icy2 regulates autophagosome formation. Autophagy 11, 2288–2299 (2015).

57. M. Dienhart, K. Pfeiffer, H. Schagger, R. A. Stuart, Formation of the yeast F1F0-ATP synthase dimeric complex does not require the ATPase inhibitor protein, Inh1. J Biol Chem 277, 39289–39295 (2002).

58. T. Hashimoto, Y. Yoshida, K. Tagawa, Purification and properties of factors in yeast mitochondria stabilizing the F1F0-ATPase-inhibitor complex. J Biochem 95, 131–136 (1984).

59. F. X. R. Sutandy, I. Gossner, G. Tascher, C. Munch, A cytosolic surveillance mechanism activates the mitochondrial UPR. Nature 618, 849–854 (2023).

60. V. K. Vyas et al., New CRISPR Mutagenesis Strategies Reveal Variation in Repair Mechanisms among Fungi. mSphere 3, (2018).

